# Drought restructures multitrophic food webs through cascading eco-physiological constraints

**DOI:** 10.64898/2025.12.19.695575

**Authors:** Bijay Subedi, Mônica F. Kersch-Becker

## Abstract

Ecological communities are linked not only by energy flows, but also by sensory and behavioral pathways that allow organisms to locate, recognize, and interact with one another. Environmental stressors such as drought may disrupt these pathways by altering both resource availability and interactions that regulate both population and community dynamics. While species-level responses to drought are well documented, far less is understood about how drought-driven changes in plant resources initiate bottom-up effects that cascade through multiple trophic levels. Here, we tested how drought affects a plant-herbivore-carnivore system by quantifying plant physiological changes and their cascading effects on herbivore performance, nutritional quality, and carnivore behavior. Drought reduced plant growth and stomatal conductance, which lowered herbivore feeding efficiency, survival, and reproduction. Herbivores feeding on drought-stressed plants also displayed lower protein content and biomass. Drought also suppressed the emission of herbivore-induced plant volatiles (HIPVs), limiting the chemical cues carnivores use to locate prey. Despite higher prey consumption rates in no-choice assays, carnivores suppressed herbivore populations less effectively under drought. At the community level, these disruptions caused disproportionate declines in arthropod abundance and diversity, particularly among carnivores, leading to trophic simplification. These findings show that drought not just reduce resource availability, it disrupts chemical cues and trophic feedbacks that maintain food web structure. By linking changes in resource limitations to community-level responses, this study demonstrates how environmental stressors can weaken top-down control and destabilize multitrophic dynamics. Predicting ecological resilience depends not only on responses but on how stress disrupts the information signals that maintain ecosystem services and function.

## 1. Introduction

Environmental stressors are major drivers of ecological change^1^, not only through direct physiological effects but also by reshaping the interactions that maintain communities^2,3^. Shifts in interaction strength, whether predation, competition, or mutualism, can accelerate population declines and elevate extinction risk, often compounding or even exceeding the effects of physiological stress alone^4–6^. This is because species do not exist in isolation, but are embedded in tightly linked ecological networks, where disruptions to even a single interaction can ripple outward, destabilizing trophic dynamics, breaking down mutualisms, or intensifying competition^7^. These cascading effects can propagate stress across multiple trophic levels, placing shifts in species interaction at the center of ecological change within communities^7,8^. Among stressors, drought is a globally pervasive and increasingly frequent selective filter, influencing not only individual physiology but also the structure and stability of entire ecological communities^9–11^. These community-level effects often emerge through disruptions in foundational trophic relationships, particularly those among plants, herbivores, and carnivores, that drive energy flow, population regulation, and ecological resilience. Understanding how environmental stress, such as drought, modifies these trophic interactions is crucial for anticipating broader shifts in community structure and dynamics.

Traditionally, the control of herbivore populations has been attributed to either bottom-up or top-down forces^12^, accumulating evidence, however, has demonstrated that these forces interact. Yet, less is known how environmental factors, such as drought, can affect both forces, which could produce context-dependent patterns of community assembly^11,13,14^. Plant mitigate the effects of drought by reallocating resources among competing physiological processes^15^. Reduced water availability suppresses the biosynthesis of key nutritional compounds, often lowering tissue quality and altering defensive chemistry^15,16^. While these changes in plant physiological effects are well established, how they scale up to higher trophic levels remains understudied. Herbivores, particularly phloem feeders such as aphids, are highly sensitive to drought-induced declines in host plant quality, where reduced turgor pressure limits feeding and often force compensatory behaviors with metabolic costs^19–21^. Such plant-mediated stressors can also interact with intraspecific processes, including density-dependent competition, shaping herbivore growth and reproduction. However, it is unclear whether drought modifies the strength or outcome of these density-dependent effects. Understanding whether these physiological and demographic constraints also diminish the nutritional value of herbivores as prey is critical for linking bottom-up stress to higher trophic responses.

Carnivores face compounded challenges^22^ as they rely on herbivore-induced plant volatiles (HIPVs) for finding prey^23,24^. Drought suppresses plant’s volatile emissions and alters volatile blend composition^25^. These changes can interfere with carnivore foraging, potentially reducing prey detection and hunting efficiency. Concurrently, reduced prey availability and quality can constrain carnivore efficacy^26^, creating mismatches between carnivore presence and regulatory function of herbivore population^27–29^. Determining how these interacting constraints weaken top-down control will help resolve a central question in stress ecology: how environmental filters destabilize trophic regulation.

At the plant-associated community level, drought can restructure multitrophic assemblage by altering the abundance and interactions of herbivore, carnivores, and other functional guilds^13,14^. This restructuring can reshape trophic architecture, disrupt energy flow, and erode functional diversity, ultimately reducing ecosystem stability and resilience. Community-wide responses often emerge from nonlinear feedbacks and interaction mismatches, making them difficult to predict from species-level traits alone. To anticipate biodiversity loss and the breakdown of ecological function under increasing environmental stress, it is essential to understand how environmental filters reorganize multispecies assemblages. Although species level physiological responses to drought are well characterized^11^, we lack a predictive framework for how they scale up to multitrophic community dynamics.

In this study, we identify the mechanisms through which drought-driven plant responses restructure arthropod assemblages, advancing beyond descriptive patterns of community organization. Through greenhouse, lab, and field experiments, we describe how drought-induced plant responses cascade through trophic levels altering herbivore performance and disproportionately affecting carnivores within the arthropod community. Specifically, we test whether (i) drought-induced declines in plant performance constrain aphid feeding efficiency and population growth; (ii) suppression of HIPVs under drought impairs carnivore visitation and reduces biological control; and (iii) these combined effects scale to reduce arthropod community diversity and produce trophic bottlenecks. This integrative approach provides a framework for predicting how environmental stress reshapes species interactions and reduces ecological stability.

## 2. Materials and Methods

### 2.1. Study system and experimental design

We used tomato plant (*Solanum lycopersicum* cv. Moneymaker) and herbivore, potato aphids (*Macrosiphum euphorbiae,* (Hemiptera: Aphididae)) to investigate how drought stress affects multitrophic interactions. The generalist predator, convergent lady beetle (*Hippodamia convergens,* (Coleoptera: Coccinellidae)), was incorporated in a subset of experiments to examine top-down effects. We conducted the experiments from May to August between 2022 and 2024, in greenhouse, laboratory, and field settings at The Pennsylvania State University, including the Russell E. Larson Agricultural Research Center (Rock Springs, PA, USA). In a full-factorial design, we manipulated two factors: water (well-watered vs. drought) and natural enemy community exposure (excluded vs. exposed). We integrated responses across biological scales, from plant physiological traits to arthropod community structure, within a unified experimental framework to identify the processes linking drought stress to multitrophic organization (Supplementary Fig. S1).

### 2.2. Plant cultivation and water treatments

Two tomato seeds were sown in 10-cm (for greenhouse experiments) and 20-cm (for field experiments) pots filled with Sunshine Mix #1 (Sungro Horticulture, USA) and grown in a greenhouse at 22□±□2□°C under a 16:8 h L:D photoperiod. Seedlings were thinned to one per pot after emergence and fertilized with 15-9-12 NPK slow-release fertilizer (Osmocote Plus, Scotts Miracle-Gro). Plants were grown for four weeks to the four-leaf stage before initiating drought treatments.

Water availability was manipulated for 5 days before any bioassay by adjusting soil volumetric moisture content: well-watered: 75–80% pot capacity, and drought: 10–15% pot capacity (Supplementary Fig. S1). Moisture content was measured every other day using a ECOWITT WH0291 Soil Moisture Tester (ECOWITT, Shenzhen, China). Watering volumes were calculated via a gravimetric calibration curve (see Supplementary Fig. S2). Larger field pots received proportionally scaled volumes based on dry soil weight relative to the standard 116 g used in 10-cm pots.

### 2.3. Effect of drought on plant traits

To assess the physiological and morphological responses of plants to drought and natural enemy treatments, we quantified four key plant traits: stomatal conductance, number of leaves, plant height, and aboveground dry biomass. These traits reflect drought-induced shifts in plant quality that can influence herbivore performance and higher trophic interactions.

We measured stomatal conductance twice per week for two weeks using a LI-600 Porometer (LI-COR Environmental, Lincoln, NE, USA) on the fourth fully expanded leaf. Stomatal conductance values were averaged across sampling dates for each individual plant, which was used as the response variable in a generalized linear mixed model (*GLMM*) assuming its gaussian distribution, fitted using the *glmmTMB* package in R 4.5.1^30^. Fixed effects included water and natural enemy exposure, with block as a random effect. The analysis was conducted on replicate-level means, comprising 124 observations across three independent trials and four treatment combinations (n=31 replicates per combination).

At the end of the experiment (i.e., Day 49), we measured plant height, total number of leaves, and aboveground dry biomass (dried at 72 °C for 48 hours and weighed using a precision balance) (n=31 replicates across three trials). For each trait, *GLMMs* with gaussian error distribution were fitted using the *glmmTMB* package, with water and natural enemy exposure as fixed effects, and block nested within trial as a random effect structure (1|trial/block). Leaf number was log-transformed to improve model fit and meet distributional assumptions. Model significance was assessed using Type II Wald chi-square tests via the *Anova ()* function from the *car* package^31^.

### 2.4. Aphid responses to drought

#### 2.4.1 Aphid survivorship and feeding efficiency (honeydew excretion)

To evaluate aphid survivorship under drought stress, we placed two adult aphids on the 3^rd^ and 4^th^ fully expanded leaves on each plant and allowed them to reproduce for 48 h. After 48 h, we removed the adult aphids and monitored the survival of newly born nymphs (n=2 nymphs/plant) daily until their death (n=30 plants replicates per treatment). A Cox proportional hazards mixed model was fitted using the *coxme* package^32^, with water treatment as fixed effect and plant ID as a random effect and visualized using Kaplan-Meier curves using *survminer* package^33^.

We estimated aphid feeding efficiency by measuring honeydew excretion^34^. Ten apterous adults were enclosed per plant using custom-made clip cages lined with pre-weighed aluminum foil cups (Supplementary Fig. S3). After 72 hours, we recorded the total number aphids in the clip-cage and weighed the cups using a precision microbalance (XP26DR Excellence Plus Microbalance, Mettler Toledo, Columbus, OH, USA). Honeydew production was quantified as the difference between the final and initial weight of each foil cup, and aphid feeding efficiency was calculated as mg honeydew per aphid. Two trials were conducted (n=10 replicates per treatment per trial, trials=2). A *GLMM* with gaussian error distribution was fitted using *glmmTMB*, with water treatment and trial as fixed effects and block as a random effect.

#### 2.4.2 Aphid population dynamics (greenhouse and field)

In the greenhouse, we introduced a mixture of late instar and adult aphids at four initial densities (5, 10, 50, and 100 individuals) onto caged plants (25 × 25 × 50 cm) (n=21 replicates across three trials). These density treatments in greenhouse experiment were designed to test whether drought alters the strength or shape of density-dependent population growth in aphids. In the field, each plant was inoculated with 50 aphids (mix of late instar and adult aphids; n=31 replicates across three trials) and placed under a rain-exclusion shelter (13 × 17 m) with a transparent polyethylene roof and open sides to allow natural arthropod colonization. We manipulated natural enemy community by either excluding them with mesh cages or allowing their access by leaving plants uncaged (Supplementary Fig. S4). We counted the number of aphids on each plant on Days 4, 9, and 14. Data from both the greenhouse and field experiments were analyzed separately using two approaches:

a. For aphid final density, repeated-measures *GLMMs* (negative binomial distribution) using *glmmTMB* with water, initial aphid density (for greenhouse), natural enemy exposure (for field), time, and their interactions as fixed effects; trial/block, and plant ID as random effects.
b. Per capita population growth rate (PGR) was computed as: (*dN/Ndt*) = ln (*N_2_*-*N_1_*)/(*t_2_*-*t_1_*), where *N_1_* and *N_2_* are initial and final aphid densities, respectively^35^. Gaussian *GLMMs* (package: *glmmTMB*) were fitted to PGR using water, natural enemy exposure (for field only), and initial aphid density (for greenhouse only) as fixed effects and trial/block as random effects.

#### 2.4.3 Aphid quality assessment

To assess how drought and natural enemy presence affect aphid physiological traits, we collected 10 adult aphids per plant from the second field trial (n=12/treatment). We estimated aphid mass by dividing the total mass by the total number of aphids. Protein content was quantified using the Bradford assay (Bradford, 1976) following homogenization in 500 μL extraction buffer. To analyze total mass and protein content, we conducted *GLMMs* (package: *glmmTMB*) with gaussian error distribution with water and natural enemy as fixed effects and block as random effects.

### 2.5. Effects of drought on natural enemy responses

#### 2.5.1 Predator effect

To evaluate how drought affects predator suppression of aphid populations in field, we estimated predator effect (consumptive and non-consumptive) by comparing observed aphid densities on natural enemy-exposed plants to predicted densities based on growth rates from exclusion cages using the formula: predicted = *I + D x P*, where *I* = initial aphid density, *D* = experiment duration (14 days), and *P* = per diem growth rate calculated from exclusion cages^36^. Predator effect = predicted – observed aphid abundance. This metric integrates both consumptive effects (direct predation) and non-consumptive effects (e.g., predator-induced reductions in aphid reproduction or dispersal) associated with the presence of natural enemies. Predator effect was analyzed using a gaussian *GLMM* (*lme4*)^37^ with water as a fixed effect and trial/block as random effects.

#### 2.5.2 No-choice consumption assay

We placed ten aphids per treatment (collected from the second field trial plants; n=12/treatment) into agar-coated Petri dishes along with a single 24 h starved adult *H. convergens*. We recorded aphid consumption at 30-, 60-, 100-, and 180-minutes. We tested the effects of drought and natural enemy exposure on predator consumption using Cox proportional hazards mixed models (package: *coxme*), with replicate nested within Petri dish as a random effect.

### 2.6. Herbivore-induced plant volatiles (HIPVs) emissions

To evaluate how drought and exposure to the natural community of arthropods affect carnivore-attractance volatile cues, we collected volatile organic compounds (VOCs) from the second and third field trials, three days after aphid inoculation from the third fully expanded leaf using a dynamic headspace trapping method (n=14/treatment across two trials)^38^. We enclosed the 3^rd^ fully expanded leaf in a 1240 mL polyethylene cups and trapped volatiles for 6 hours using HayeSep-Q filters (Supelco, Bellefonte, PA, USA)^39^. We eluted the compounds with 200 μL dichloromethane containing 10 ng/μL Nonanyl acetate as internal standard and analyzed on a Gas Chromatography–Mass Spectrometry (GC-MS) (Agilent 7890A/5975C). We first removed low-confidence and contaminant compounds prior to analysis, retaining only high-quality volatile detections (match score ≥ 80). Compound abundances were normalized to an internal standard, nonanyl acetate (2 ng/uL). To assess treatment effects on overall VOC emissions, we used *GLMM* with zero-inflated gamma error (package: *glmmTMB*) selecting the best fit using *AIC*. Differences in total emission across treatment were tested using *ANOVA* (package: *car*) and post hoc comparisons were performed using Tukey-adjusted means. To evaluate changes in VOC composition, we conducted permutational multivariate analysis of variance (*PERMANOVA*) using Bray–Curtis dissimilarities, followed by pairwise comparisons between treatment groups (package: *vegan*). Assumptions were checked via homogeneity of dispersions (*PERMDISP*), and differences were visualized using non-metric multidimensional scaling (*NMDS*). Vector fitting (*envfit*) was used to identify key compounds contributing to observed shifts. We further analyzed a subset of 16 ecologically relevant compounds using multivariate analysis of variance (*MANOVA*), followed by univariate *ANOVAs* and post-hoc with *tukey* adjustment.

### 2.7. Arthropod community structure and composition in response to drought

We conducted visual arthropod census twice per week for two weeks on natural enemy-exposed plants to assess how drought stress influences arthropod community composition and trophic interactions. We identified individuals to the lowest feasible taxonomic level and assigned them to feeding guilds: carnivore (including predators and parasitoids), omnivores, pollinators, herbivore (including chewers and suckers), and scavengers. To characterize community composition, we aggregated individuals by feeding guild.

We used Canonical Correspondence Analysis (CCA) to assess the influence of drought on arthropod community composition. Abundance data were Hellinger-transformed to reduce the influence of rare species and meet assumptions of unimodal species responses. To account for the experimental design, block nested within trial was included as a conditional term in the model. Model significance was assessed using permutation-based ANOVA (999 permutations) for the full model and individual predictor terms. Species associations with the ordination axes were evaluated using vector fitting (*envfit*), with significance determined via 999 permutations.

To assess how drought influenced arthropod community diversity, we examined three ecological response metrics, abundance, species richness, and Shannon-Wiener (H′) diversity, for each functional guild and the overall arthropod community. We first conducted a *MANOVA* using Roy’s greatest root to detect overall multivariate effects of drought. When this test revealed significant differences, we followed up with univariate models for each metric^40^. Specifically, we used generalized linear mixed models (*GLMMs*) with a negative binomial error structure for abundance and richness, which are discrete and overdispersed variables. For Shannon diversity, we applied a Gaussian linear model appropriate for continuous, normally distributed data. In all univariate models, we included water, arthropod feeding guild, and their interaction as fixed effects, and incorporated block as a random effect to account for spatial variability in arthropod communities across the experimental field.

## 3. Results

### 3.1 Effect of drought on plant traits

Drought stress substantially impaired plant physiological and morphological performance across all measured traits. Drought-stressed plants produced ∼38% fewer leaves (χ²_(1)_=48.30, *P*<0.001) compared to well-watered plants, with leaf number declining from 15 to 11 on average. (Supplementary Fig. S5a). Additionally, drought-stressed plants were ∼55% shorter than well-watered plants, averaging 21 cm in height difference (χ^2^_(1)_=229.09, *P*<0.001; Supplementary Fig. S5b).

Aboveground dry biomass was particularly sensitive to drought, decreasing by ∼140%, with mean biomass falling from ∼31.0 g in well-watered plants to ∼13.0 g under drought (χ^2^_(1)_=481.32, *P*<0.001; Supplementary Fig. S5c). Stomatal conductance exhibited the most pronounced decline, decreasing by over 340%, with drought-stressed plants showing mean values of ∼0.117 mol m□² s□¹ compared to ∼0.534 mol m□² s□¹ in well-watered plants (χ^2^_(1)_=1672.02, *P*<0.001; Supplementary Fig. S5d). In contrast, natural enemy or drought-by natural enemy interaction exposure had no statistically significant effect on any plant trait measured (all *P*-value>0.05; Supplementary table S1 & Fig. S5). These results indicate that drought, rather than trophic interactions, was the dominant driver of plant performance in this system.

### 3.2. Aphid responses to drought

#### 3.2.1 Aphid survivorship and feeding (honeydew excretion)

Drought stress shortened aphid lifespan by eight days compared to the well-watered treatment (Log rank test, global comparisons: χ^2^_(1)_=125, *P*<0.0001; Fig. 1a). Drought reduced aphid honeydew excretion (χ^2^_(1)_=31.79, *P*<0.001), where aphids feeding on drought-stressed plants excreted 61% less honeydew per capita compared to those feeding on well-watered plants (Fig. 1b).

**Fig. 1.**
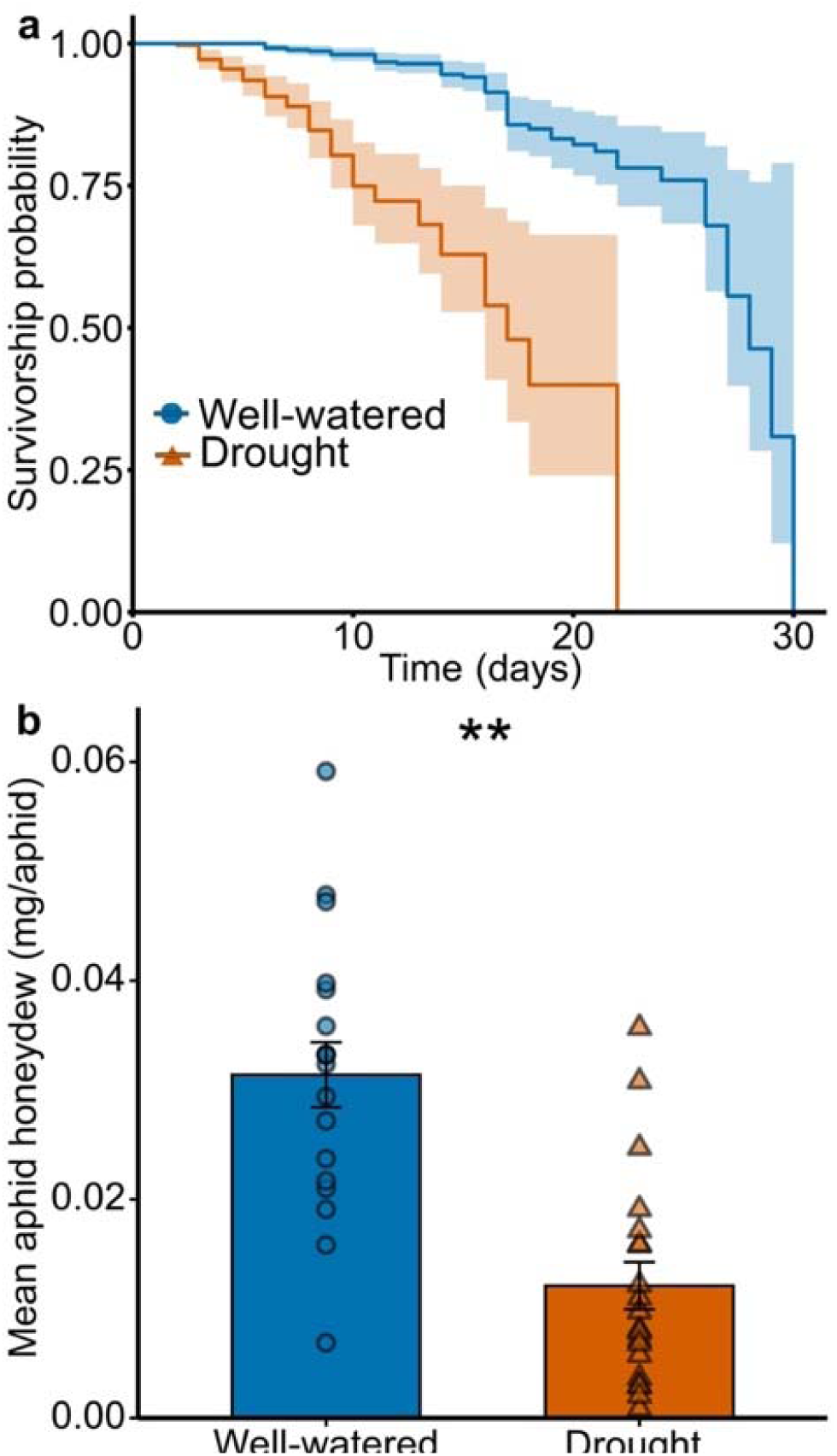
Effects of drought on aphid survival and honeydew production. (a) Kaplan–Meier survivorship curves of aphids on well-watered and drought-stressed plants, with shaded areas representing 95% confidence intervals. (b) Mean aphid honeydew excretion (mg per aphid; mean ± SE) on well-watered and drought-stressed plants. Individual data points are shown, and asterisks indicate significant differences between treatments (*P*<0.01).

#### 3.2.2 Aphid population dynamics (greenhouse and field)

##### 3.2.2.1 Density dependence experiment

Per capita aphid growth rate (PGR) was significantly influenced by an interaction between drought and aphid density (χ^2^_(3)_=35.14, *P*<0.001; Fig. 2a). On well-watered plants, PGR declined as initial density increased, consistent with negative density-dependent population growth under resource-sufficient conditions. In contrast, aphid populations on drought stressed plants show suppressed growth even at low initial densities, suggesting density-independent regulation, likely due to persistent host resource limitation.

**Fig. 2.**
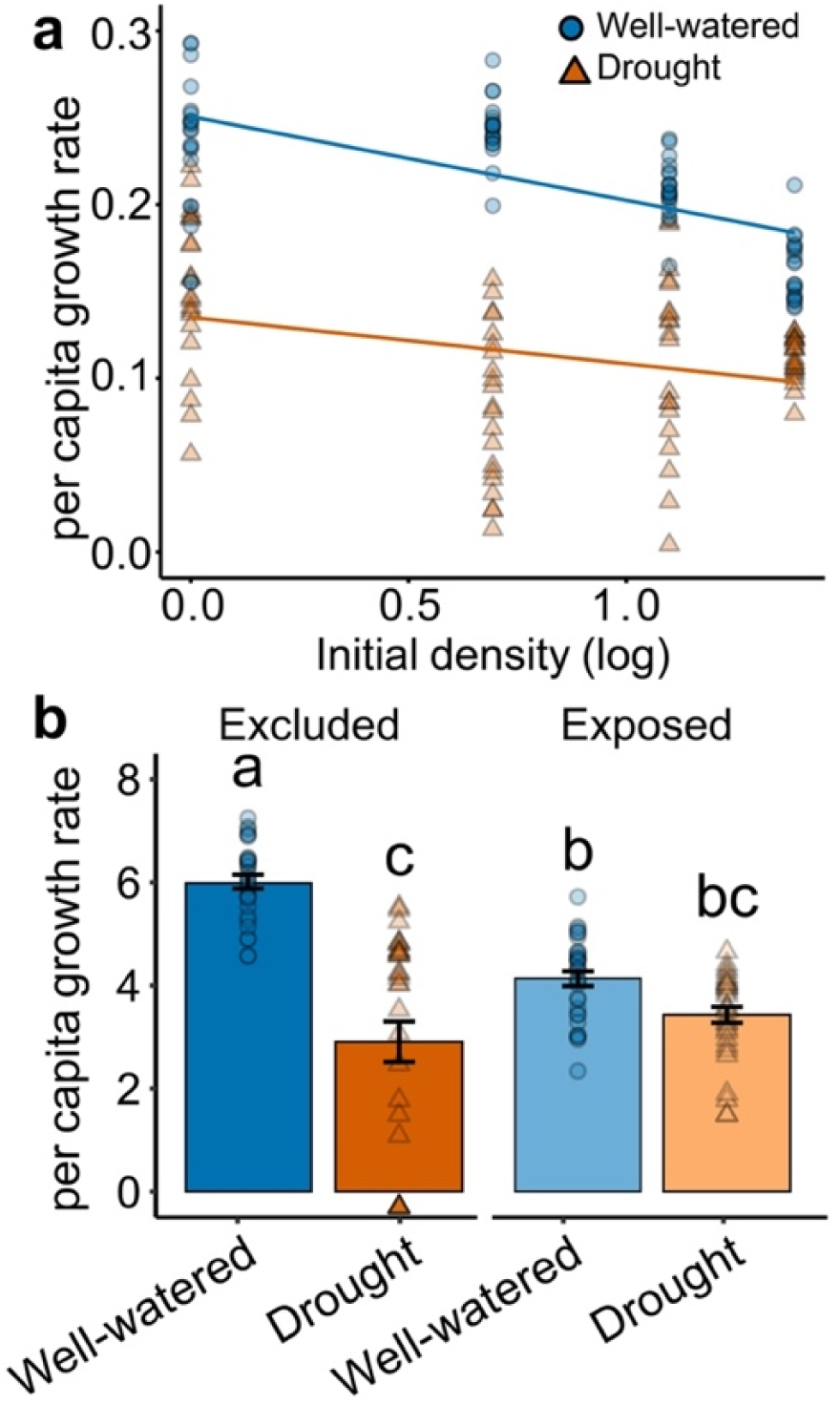
Effects of drought stress on *Macrosiphum euphorbiae* per capita population growth rate in response to initial aphid density and drought stress in the greenhouse (a), and in response to water and natural enemy exposure treatment in field (b). Blue line, bars, and symbols represent well-watered, and orange line, bars, and symbols represent drought conditions.

A significant three-way interaction among drought, aphid density, and time also influenced total aphid abundance over time (χ^2^_(3)_=17.14, *P*<0.001; Supplementary Fig. S6). Well-watered plants supported consistently higher aphid densities across all time points, with final aphid abundances ranging from 220% to 440% greater than drought-stressed plants depending on initial aphid density (Supplementary Fig. S6). These differences increased with time and density, consistent with a compounding effect of favorable host resources on population growth.

##### 3.2.2.2 Field Experiment

Similarly, in the field, the strength of this top-down suppression varied by water availability (drought × natural enemy exclusion: χ^2^_(1)_=41.98, *P*<0.001; Fig. 2b). In well-watered plants, carnivore presence reduced aphid PGRs by ∼31%. In contrast, under drought, the suppressive effect was far weaker: natural enemies reduced PGRs by only ∼15%. These differences highlight that predators exert stronger control when aphids feed on well-watered hosts. Aphid per capita population growth rate was also affected by the water × natural enemy × time interaction. On well-watered plants, PGRs were more than double those under drought in enemy-exclusion cages (χ^2^_(1)_=103.22, *P*<0.001; Fig. 2b).

Aphid population processes were shaped by an interactive effect of drought and natural enemy exclusion. By Day 14, aphid abundance on well-watered plants was over 900% higher than on drought-stressed plants under natural enemy exclusion, but only 110% higher when natural enemy were present, indicating that bottom-up reduced top-down effectiveness (drought × natural enemy interaction: χ^2^_(2)_=62.61, *P*<0.001; drought × time: χ^2^_(2)_=90.07, *P*<0.001; drought × natural enemy × time: χ^2^_(2)_=32.51, *P*<0.001; Supplementary Fig. S7). These results indicate that both trophic and abiotic constraints jointly determine population-level outcomes under fluctuating environmental conditions.

#### 3.2.3 Aphid quality

Drought significantly reduced aphid mass, with aphid mass from drought-stressed plants, weighing 47% less (χ^2^_(1)_=146.95, *P*<0.001; Fig. 3a) compared to aphids from well-watered plants. The presence of carnivores affected aphid mass, aphid exposed to the natural enemy community weighed 26% less than aphids within enclosures (χ²=16.69, df=1, *P*<0.001; Fig. 3a). However, the interaction between drought and natural enemy exposure did not significantly affect aphid mass (χ^2^_(1)_=1.465, *P*=0.226; Fig. 3a).

**Fig. 3.**
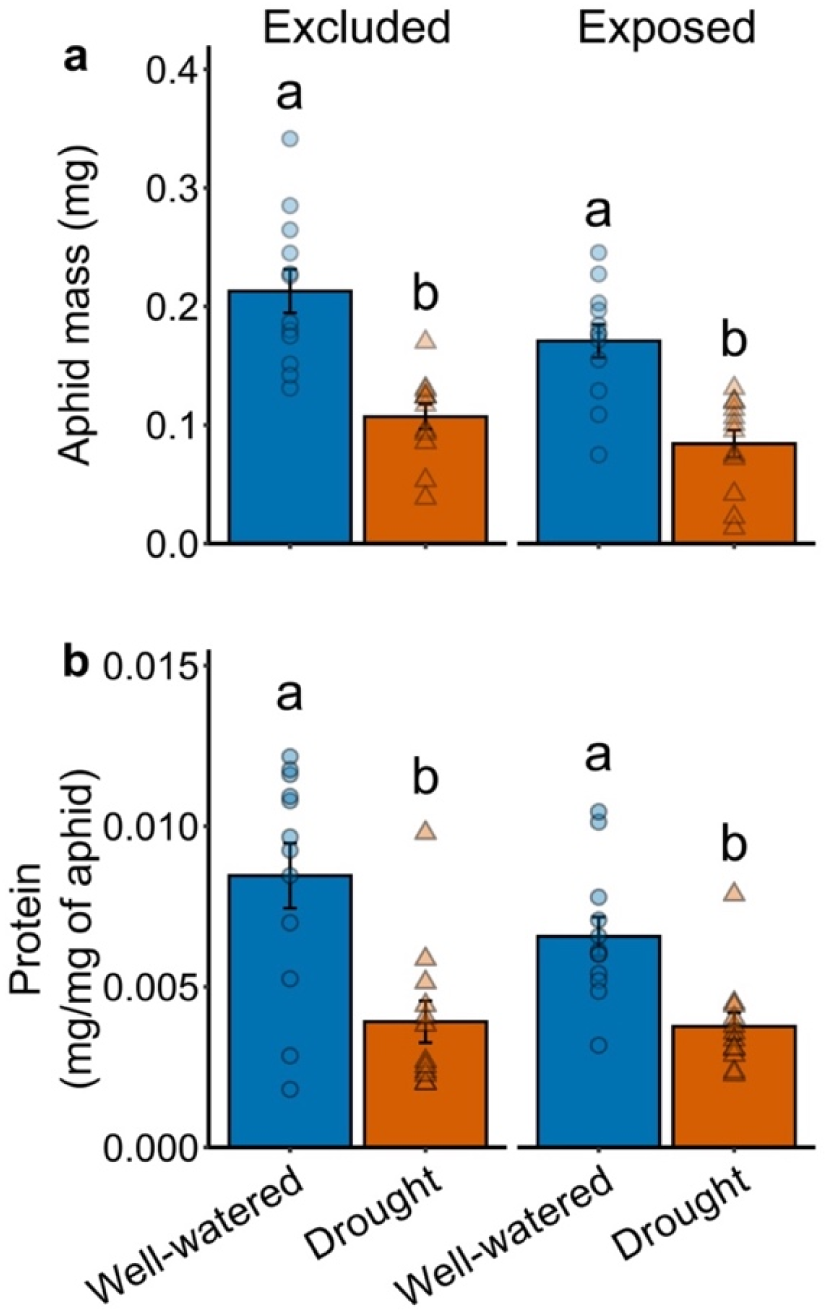
Aphid mass (a), and protein content per milligram, (b) of aphid under well-watered and drought treatment and natural enemy excluded or exposed (*P*<0.05).

Drought also strongly reduced aphid protein content. Aphids feeding on drought-stressed plants had 45% lower protein content (χ^2^_(1)_=29.853, *P*<0.001; Fig. 3b) compared to under well-watered plants. Neither the natural enemy exposure nor its interaction with water treatment significantly influenced aphid protein content (natural enemy: χ^2^_(1)_=2.297, *P*=0.129; Fig. 3b; interaction: χ^2^_(1)_=1.688, *P*=0.194; Fig. 3b).

### 3.3 Natural enemy responses

#### 3.3.1 Predator effect

In field conditions, when natural enemies can choose whether to attack aphid on drought-stressed or well-watered plants, the estimated predation rate was 24 times higher in well-watered plants compared to the drought-stressed plants (χ^2^_(1)_=53.14, *P*<0.001; Fig. 4a). The number of aphids present at the final count, included as a covariate, did not significantly influence predation rate (χ^2^_(1)_=0.495, *P*=0.48).

**Fig. 4.**
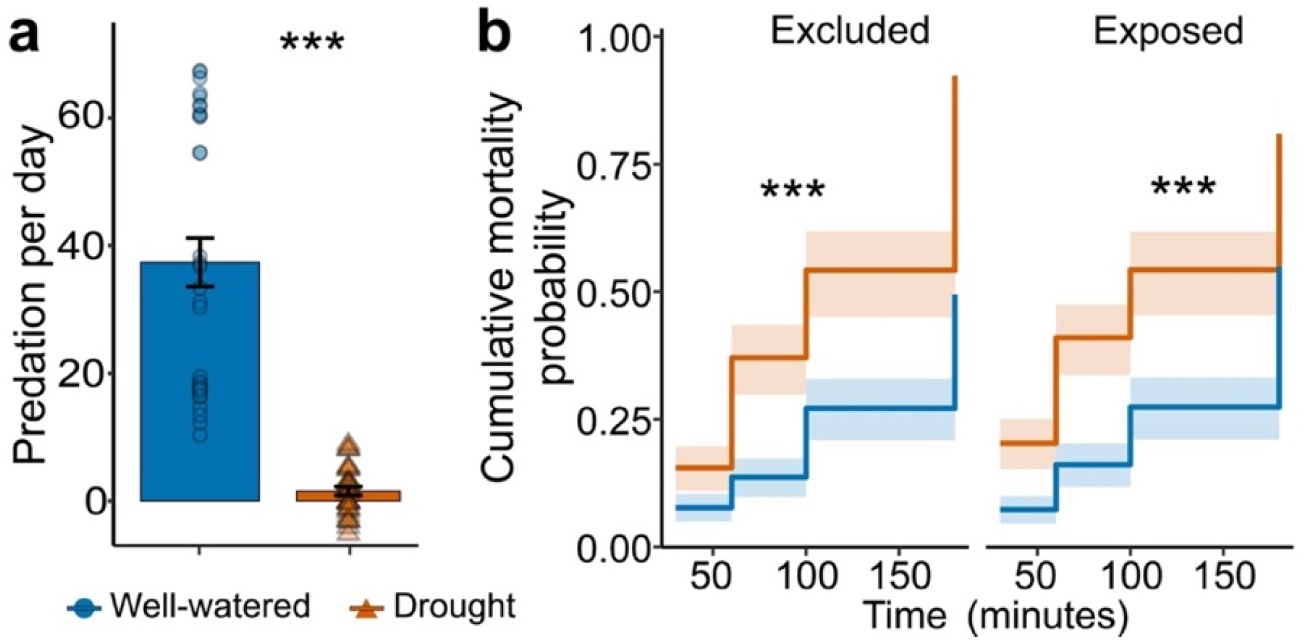
Effects of drought and exposure to natural enemies on (a) field estimates of predation rate (P< 0.05), and (b) ladybug predation, presented as cumulative mortality probability of aphids, in no-choice petri-dish consumption assay using aphids collected from the field experiment.

#### 3.3.2 No-choice consumption assay

In no-choice experiments, drought significantly increased predation by 405% (Z=-5.30, *P*<0.001; Fig. 4b). Previous exposure to natural enemies did not affect aphid predation (Z=0.64, *P*=0.525; Fig. 4b), and the interaction between drought and natural enemy treatment was also not significant (Z=-0.14, *P*=0.886; Fig. 4b).

### 3.4. Herbivore-induced plant volatiles (HIPVs) emissions

Drought significantly reduced total volatile organic compounds (VOC) emissions (χ^2^_(1)_=19.45, *P*<0.001; Fig. 5a), regardless of natural enemy exposure. Compared to well-watered plants, drought reduced total VOC emissions by 85%. Natural enemy exposure alone did not affect total emissions and did not interact with drought (natural enemy: χ^2^_(1)_=1.232, *P=*0.267; interaction: χ^2^_(1)_=0.334, *P=*0.563; Fig. 5a). These results show that drought alone strongly suppressed overall volatile emissions.

**Fig. 5.**
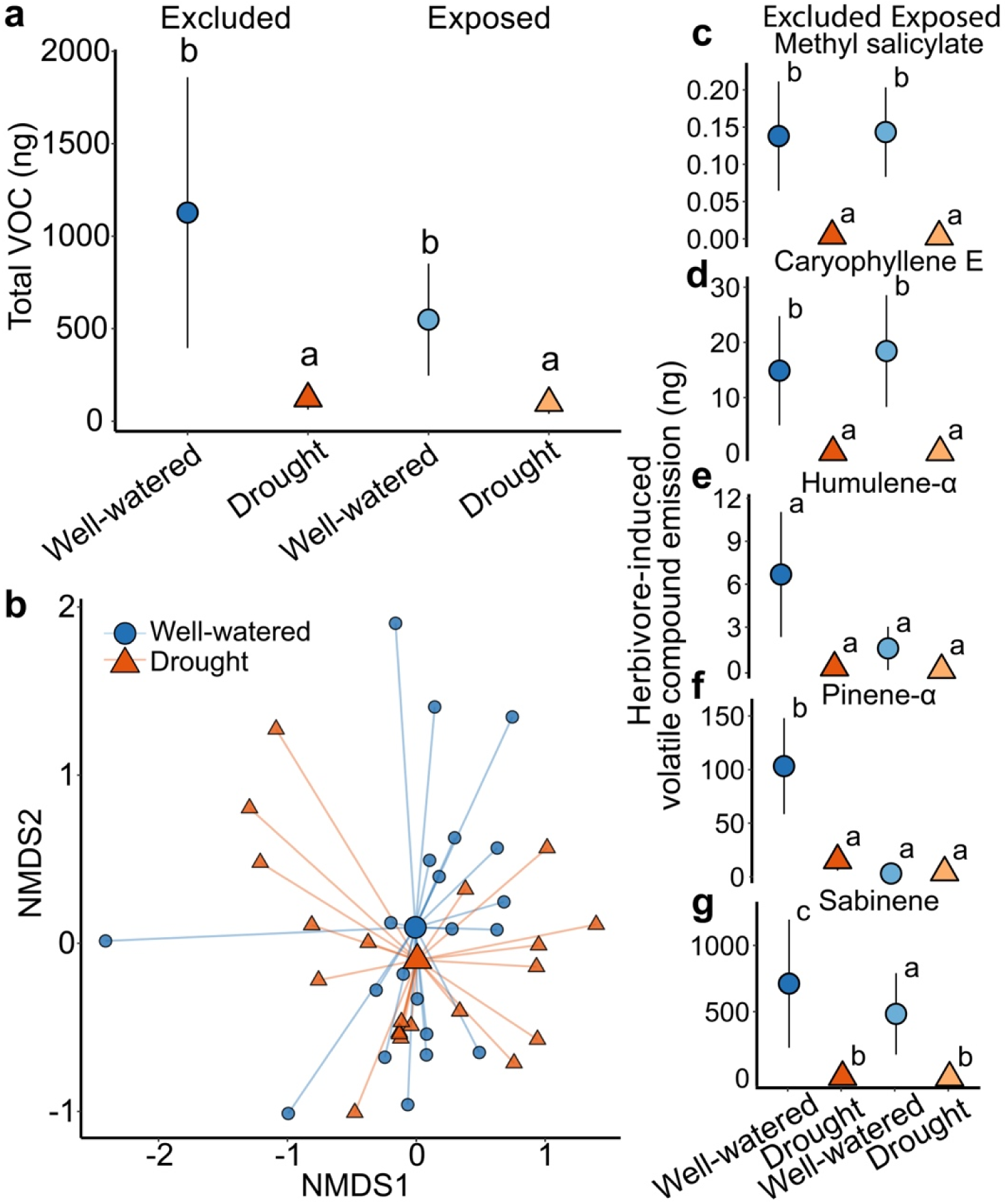
Effects of drought and natural enemy exposure on herbivore-induced plant volatile (HIPV) emissions and composition. (a) Total HIPV emissions (mean ± SE) across four treatment combinations: well-watered and drought-stressed plants either exposed to or excluded from natural enemies. Groups with different letters indicate significant differences based on Tukey’s HSD tests (*P* < 0.05). (b) Non-metric multidimensional scaling (NMDS) ordination of HIPV profiles from tomato plants, with each point representing an individual plant (PERMANOVA, *P*<0.05). Centroids summarize group-level volatile profiles. (c–g) Emissions of selected individual compounds (mean ± SE) across treatments, with significant differences among groups indicated by different letters (*P* < 0.05).

Multivariate analysis showed that drought significantly affected VOC blend composition, whereas natural enemy exposure did not. Using *PERMANOVA* with Bray–Curtis dissimilarity, we detected a significant drought effect (F_(1,38)_=4.024, *P*=0.001; Fig. 5b) but found no significant effect of natural enemy exposure (F_(1,38)_=1.19, *P*=0.288) or the interaction (F_(1,37)_=0.95, *P*=0.47). In the compound level vector fitting (*envfit*), key compounds driving the observed shift in HIPV composition were methyl salicylate (MeSA), cumin aldehyde, nonanal, and decanal. Pairwise post-hoc comparisons among treatment combinations revealed several significant differences in VOC profiles. Drought-stressed plants exposed to natural enemies exhibited significantly different VOC profiles compared to well-watered plants exposed to natural enemies (F_(1,17)_=2.69, *P*=0.004; Supplementary Fig. S8) and well-watered plants protected from natural enemies (F_(1,20)_=2.28, *P*=0.039; Supplementary Fig. S8). We found no significant differences between natural enemy exposed and natural enemy protected plants within either water treatment (drought: F_(1,18)_=1.009, *P*=0.386; well-watered: F_(1,19)_=1.13, *P*=0.333; Supplementary Fig. S8), suggesting that natural enemy exposure alone did not substantially alter VOC profiles within a given water treatment.

MANOVA indicated significant effects of drought (Pillai’s trace=0.683, F_(16,22)_=2.964, *P*=0.009), but not of natural enemy exposure (Pillai’s trace=0.573, F_(16,22)_=1.84, *P*=0.093) or interaction (Pillai’s trace=0.522, F_(16,22)_=1.50, *P*=0.185). Drought significantly suppressed emissions of several HIPVs known to be carnivore attractants. It reduced MeSA emissions by 97% (F_(1,19)_=8.27, *P*=0.007; Fig. 5c), E-Caryophyllene by 99% (F_(1,19)_=5.20, *P*=0.028; Fig. 5d), and α-Humulene by 97% (F_(1,19)_=3.00, *P*=0.092; Fig. 5e). It also decreased α-Pinene emissions by 85% (F_(1,19)_=3.27, *P*=0.079; Fig. 5f). Sabinene emissions were completely suppressed under drought condition, with a marginal treatment effect (F_(1,19)_=4.02, *P*=0.052; Fig. 5g). No significant drought effects were detected for the remaining compounds (all *P*>0.05; Supplementary Fig. S9 & Table S3).

### 3.5. Arthropod community responses

Drought significantly reduced arthropod abundance, species richness, and Shannon-Wiener (H’) diversity, as indicated by the *MANOVA* (Roy’s greatest root=0.093, approximate F_(3,94)_=10.15, *P*<0.001). Follow-up univariate analyses showed that drought reduced arthropod richness by 47% (χ^2^_(1)_=28.01, *P*<0.001; Fig. 6a), abundance by 56% (χ^2^_(1)_=32.17, *P*<0.001; Fig. 6b), and Shannon diversity by 57% (χ^2^_(1)_=27.08, *P*<0.001; Fig. 6c) compared to well-watered plants.

**Fig. 6.**
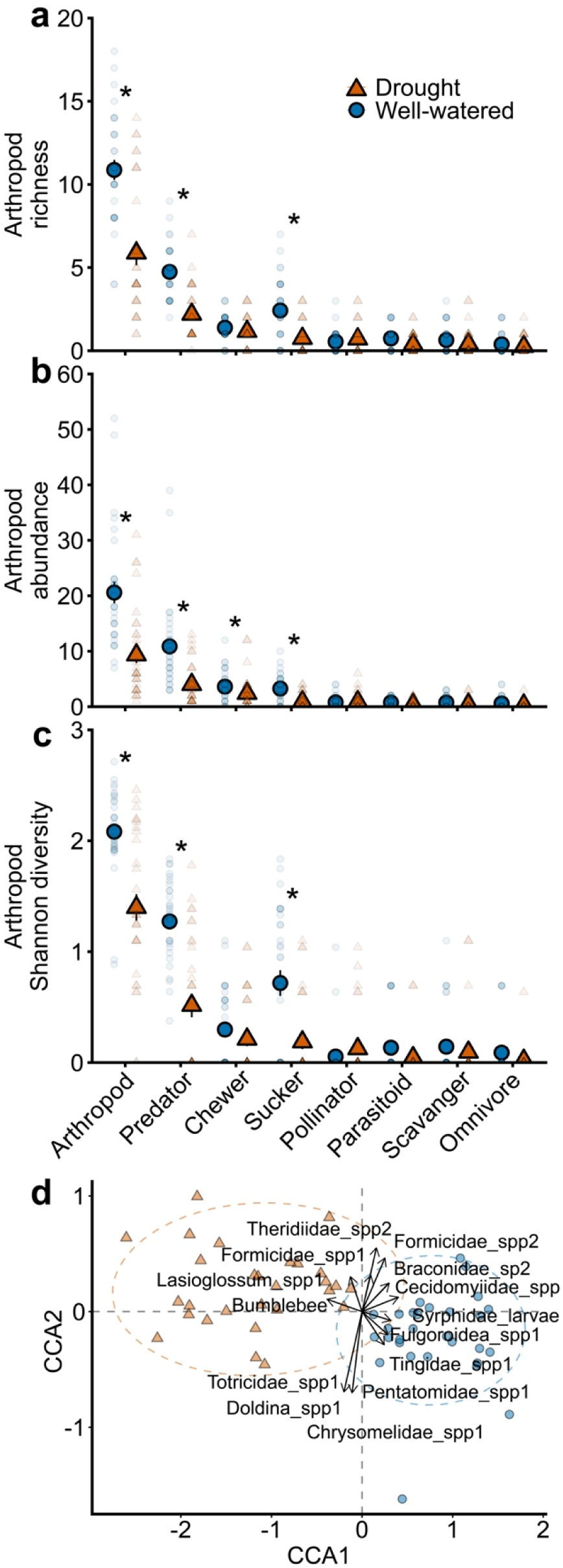
Effects of drought on arthropod community structure and composition. Effects of drought on arthropod community structure and composition. (a) Arthropod richness, (b) arthropod abundance, and (c) Shannon diversity across functional feeding guilds under well-watered (blue bars and circles) and drought (orange bars and triangles) conditions. Point represent means ± SE; individual symbols represent raw data points per plant; asterisks denote statistically significant differences (*P*<0.05) based on pairwise comparisons conducted within each trophic guild. Arthropod metrics are grouped by guild and ranked approximately by mean magnitude, with total arthropod community (“arthropod”) shown first in each panel. (d) Canonical correspondence analysis (CCA) ordination of arthropod community composition based on species-level abundance data. 95% confidence ellipses show water treatments (well-watered (blue circles) and drought (orange triangles)). Vectors represent taxa most strongly associated with treatment separation.

Drought reduced chewing herbivore abundance by 33.9% (χ^2^_(1)_=7.65, *P*=0.006; Fig. 6b). However, its effects on chewer richness and diversity were not statistically significant (P>0.05; Supplementary Table S2). Drought had particularly strong effects on carnivores, reducing predator abundance by 63.2% (χ^2^_(1)_=90.61, *P*<0.001; Fig. 6b), richness by 53.74% (χ^2^_(1)_=27.63, *P*<0.001; Fig. 6a), and diversity by 59.5% (χ^2^_(1)_=40.098, *P*<0.001; Fig. 6c). Among parasitoid metrics, only richness showed a near-significant decline (χ^2^_(1)_=3.34, *P*=0.068; Fig. 6a), while changes in abundance and diversity were not (*P*>0.05; Supplementary Table S2). Drought had no statistically significant effects on omnivore, non-aphid sucker, and pollinator abundance, richness, or diversity (all *P*>0.05; Supplementary Table S2).

Insect community composition differed significantly between water treatments. The constrained CCA model was significant (F_(1,30)_=1.37, *P=*0.035; Fig. 6d), indicating that water availability structured overall community composition, and the first canonical axis captured a significant portion of community variation (F_(1,30)_=1.37, *P=*0.022), reflecting a strong gradient in species composition between drought and well-watered conditions.

Ordination revealed clear separation between communities in drought and well-watered plots (Fig. 6d). Several species showed significant associations with treatment conditions. Taxa more abundant in drought treatments included: Tortricidae (Leafroller moth, r^2^=0.52, *P=*0.001), Reduviidae (*Doldina* spp, r^2^=0.52, *P=*0.001), Chrysomelidae (Flea beetle, r^2^=0.51, *P=*0.001), Halictidae (*Lasioglossum* spp, r^2^=0.11, *P=*0.026), and Bumblebee (r^2^=0.16, *P=*0.010). In contrast, species associated with well-watered conditions included Formicidae (sp. 1, r^2^=0.108, *P=*0.039), Formicidae (sp. 2, r^2^=0.280, *P=*0.003), aphid midge adult (r^2^=0.145, *P=*0.009), Theridiidae (r^2^=0.330, *P=*0.002), Syrphidae (Syrphid fly larvae, r^2^=0.178, *P=*0.001), Fulgoroidea (r^2^=0.122, *P=*0.020), Pentatomidae (r^2^=0.143, *P=*0.014), and Tingidae (r^2^=0.108, *P=*0.020).

## 4. Discussion

Biotic interactions are not fixed attributes of ecological communities but dynamic processes whose strength, direction, and outcomes shift with environmental conditions^41,42^. Here, we investigated how drought alters the flow of energy and information across trophic levels by influencing plant physiology, herbivore traits, and the strength of carnivore-prey interactions. Drought reduced aphid performance and diminished the effectiveness of natural enemies to regulate herbivore populations. Predation rates were higher on well-watered plants despite increased consumption of drought-stressed aphids in no-choice assays. This discrepancy between choice (field results) and no-choice (laboratory results) underscores a critical distinction: ecological impact is not solely determined by carnivore consumption capacity but by prey quality and chemical information mediating foraging within field settings^43,44^. Drought-induced changes in aphid quality, such as reduced nutritional content and mass, may have reduced predator attractiveness, contributing to the weaker top-down control observed in the field. In addition, reduced predator efficacy in the field likely reflects drought driven changes in HIPVs, which acted as an “information filter” by impairing volatile-mediated predator orientation and weakening top-down regulation. This apparent paradox illustrates a disruption in the interplay between bottom-up and top-down processes. Drought altered plant traits and weakened HIPV signaling, reducing prey quality and reshaping community composition, which together diminished natural enemy effectiveness despite concurrent declines in herbivore performance. Together, these findings reveal previously unrecognized pathways through which abiotic stress destabilizes multitrophic interactions beyond what can be inferred from abundance-based metrics alone.

### 4.1 Bottom-up constraints: plant performance determines herbivore response

Drought impaired plant physiological performance, leading to reduced stomatal conductance and biomass. Since stomata regulate both CO_2_ influx^45^ and volatile emission^39^, their restricted opening under drought imposes dual constraints on ecological interactions. First, the reduced conductance limits photosynthetic carbon assimilation^46^, suppressing primary productivity and the energetic foundation supporting herbivore populations. Second, closed stomata restricts the release of HIPVs^47^, weakening signaling foraging cues used by carnivores and potentially altering herbivore host selection and dispersal. These constraints, on energy flow and trophic signaling, may converge to limit herbivore performance. Aphid survival declined under drought, alongside reduced honeydew excretion, a proxy for phloem ingestion and feeding efficiency. These individual-level responses scaled up to population-level effects, with aphids negative density-dependent population growth shifting to density-independent regulation under host plant stress^51^. In addition, aphids exposed to carnivores showed reduced mass compared to aphids protected from carnivores, suggesting that release from predation risk also contributed to trait-level variation in performance. A reduction in protein content further confirmed a decline in nutritional value of aphids as prey, consistent with previous reports of drought-induced reductions in plant amino acid and nitrogen level^20,21^. While these results support the *Resource Concentration Hypothesis*^52^, they highlight that under intensified environmental stress, herbivore demography and population dynamics are governed primarily by resource scarcity rather than density dependence^51^. At the community level, drought-led reduced primary productivity can simplify trophic networks and weak consumer-resource coupling, thereby potentially diminishing food web stability and resilience^53^. By simultaneously limiting both the energy available and the reliability of information within ecological networks, drought has the potential to destabilize multitrophic interactions, impair ecosystem functioning, and erode the capacity of communities to buffer environmental change^53,54^.

### 4.2 Top-down weakening: drought impairs natural enemy behavior and efficacy

Our results indicate that drought-mediated bottom-up effects interact with top-down processes to shape the strength of natural enemy control of aphid populations. Although carnivore consumed more aphids from drought-stressed plants under no-choice assays, natural enemy consumption in the field was substantially higher on well-watered plants where choice was possible, indicating that prey-finding cues and carnivore recruitment, rather than per capita consumption efficiency, primarily determine effective biological control.

Plant volatile analyses provide a mechanistic explanation for changes in natural enemy effect in field conditions. Drought strongly suppressed total VOCs emission and altered blend composition, with several carnivore-attractant volatile compounds^25,55^ reduced: methyl salicylate (-97%), (E)-caryophyllene (-99%), α-humulene (-97%), α-pinene (-85%), and sabinene (fully suppressed). This drought-induced “*information bottleneck*” likely weakens effective biological control^56^ and further exacerbated by reductions in prey quality that impair carnivore performance. Similar disruptions have been observed in sugar beet, where drought-stressed plants emit fewer herbivore-induced volatiles, leading to reduced attraction of parasitic wasps such as *Aphidius colemani*^57^. If such constraints persist or intensify under environmental stress regimes, they may undermine the reliability of chemical cues that stabilize carnivore-prey dynamics, especially in systems where carnivores rely heavily on HIPVs to locate prey. Future work should test whether these disruptions scale nonlinearly with drought severity.

Although carnivores consumed more aphids from drought-stressed plants under no-choice conditions, this compensatory feeding did not translate to effective control of aphid populations in the field, where the chemical landscape is more complex and volatile cues are diluted, masked, or spatially inconsistent^58,59^. However, because well-watered and drought-stressed plants were spatially interspersed, the background chemical environment was shared across treatments, suggesting that cue dilution or masking alone is unlikely to explain the observed natural enemy avoidance. The discrepancy between field (where choice plays a role) and laboratory (where natural enemies had no choice between treatments) highlights that predator efficiency in simplified assays cannot be directly scale up to ecologically realistic environments, where signaling cues and signal-to-noise ratios are substantially reduced. In such contexts, environmental stress can impair the transmission and perception of key chemical and visual cues that guide foraging and prey detection. This indicates that sensory disruption is a functional consequence of physiological stress. Thus, drought acts not only as a physiological constraint but also as a sensory filter that disrupts behavioral pathways critical to food web dynamics. Such disruption may facilitate herbivore escape^60^, weaken carnivore-prey interactions^57,61^, and reduce network stability^62^, highlighting that stress does not merely reduce interaction rates uniformly, but can rewire the structure and strength of trophic linkages^8^. The inability of carnivores to locate prey in drought-stressed environments reveals a sensory-driven breakdown in multitrophic connectivity, underscoring the need to integrate sensory ecology within food web frameworks under environmental stress.

### 4.3 Trophic bottlenecks and community simplification

Drought sharply reduced arthropod abundance, richness, and diversity, driving a clear separation of community composition between well-watered and drought treatments. Responses to drought were highly uneven across trophic guilds, leading to asymmetries in energy transfer and emergent trophic bottlenecks. Carnivores (i.e., predators and parasitoids) were disproportionately negatively affected by drought stress, while scavenger taxa were more prevalent in drought-stressed plants. Drought-driven declines in species richness and abundance reduced redundancy in carnivore assemblages, one of properties that underpins community resilience^63^. As redundancy erodes, communities become increasingly susceptible to further disturbances, reducing their ability to maintain effective biological control.

By contrast, herbivorous feeding guilds showed contrasting results. Chewing herbivores declined in abundance but showed no significant changes in richness or diversity, in contrast non-aphid suckers were not affected by drought stress. Pollinator abundance increased by 63%, but this feeding guild was mostly represented by syrphid adults hovering on plants. However, their carnivorous larvae were more frequently recorded on well-watered plants, highlighting a mismatch between pollinating adults and carnivorous larvae under drought. As a result, energy transfer through the food web was constrained, with energy accumulating at the second trophic level rather than propagating to higher trophic levels where carnivores no longer respond to prey availability

### 4.4 Conceptual implications for stress ecology

Our findings reveal that drought destabilizes ecological processes through interconnected physiological, behavioral, and community-level processes rather than isolated pairwise effects. We propose that drought functions as a multitrophic bottleneck that deconstructs the scaffolding of interaction networks through three interlinked mechanisms: (a) suppression of plant performance and nutritional quality; (b) reduced herbivore feeding, fitness, and population growth; and (c) impaired top-down regulation arising from the loss of carnivore-attractant cues and declining prey quality^13,14^. These mechanisms operate simultaneously, generating systemic rather than additive effects. By disrupting both resource flow (e.g., plant biomass) and information flow (e.g., HIPVs) that sustain trophic regulation, drought weakens the connections among trophic levels^3,64^. The result is a simplified and less resilient food web, where species persistence and functional performance decline, redundancy within trophic guilds erodes, and overall regulatory capacity deteriorates.

The novelty of our framework lies in integrating physiological, behavioral, and community-level mechanisms into a unified understanding of how abiotic stress propagates through ecological networks. Future research should integrate trait-based community ecology with network analyses of interactions to determine whether chronic stress leads to irreversible network collapse or whether systems can rebound under pulsed disturbances. Experimental approaches should also incorporate spatial and behavioral complexity, including microhabitat heterogeneity and carnivore foraging plasticity, to simulate real ecological conditions. From a predictive standpoint, models of food web resilience should incorporate not only biomass and population metrics but also informational traits such as carnivore-attractant volatiles and prey quality. In agricultural systems, integrating these variables will improve early-warning systems for biological control failure and guide targeted actions to sustain ecosystem services. More broadly, integrating informational and behavioral dimensions into community ecology and resilience frameworks not only describes how environmental stress reshapes interaction networks but also identifies strategic points for interventions to maintain or restore ecosystem stability.

## DATA ACCESSIBILITY

Data used in this manuscript are available from the Dryad Digital Repository: http://datadryad.org/share/LINK_NOT_FOR_PUBLICATION/LSwDcAxd7wyMX8i9wqjg9JAPvDmg1HIhKoJQSyRXyaE (Subedi et al. 2025).

## STATEMENT OF AUTHORSHIP

BS and MFKB conceptualized the research; BS conducted the experiments and collected the data; BS and MFKB analyzed the data. BS and MFKB wrote the paper, and all authors contributed substantially to revisions.

## ACKNOWLEDGEMENTS

We thank Erin Gall and Allyson Suman for their help with setting up the experiment and data collection. This work was supported by the Department of Entomology at the Pennsylvania State University, the National Science Foundation, Division of Environmental Biology award #2440876 the USDA National Institute of Food and Agriculture and Hatch Appropriations under Project #PEN04923, and Accession #7006440, and USDA multi-state project #PEN04757.

