## Supplementary table S1 for "Drought restructures multitrophic food webs through cascading eco-physiological constraints"

**Figures**

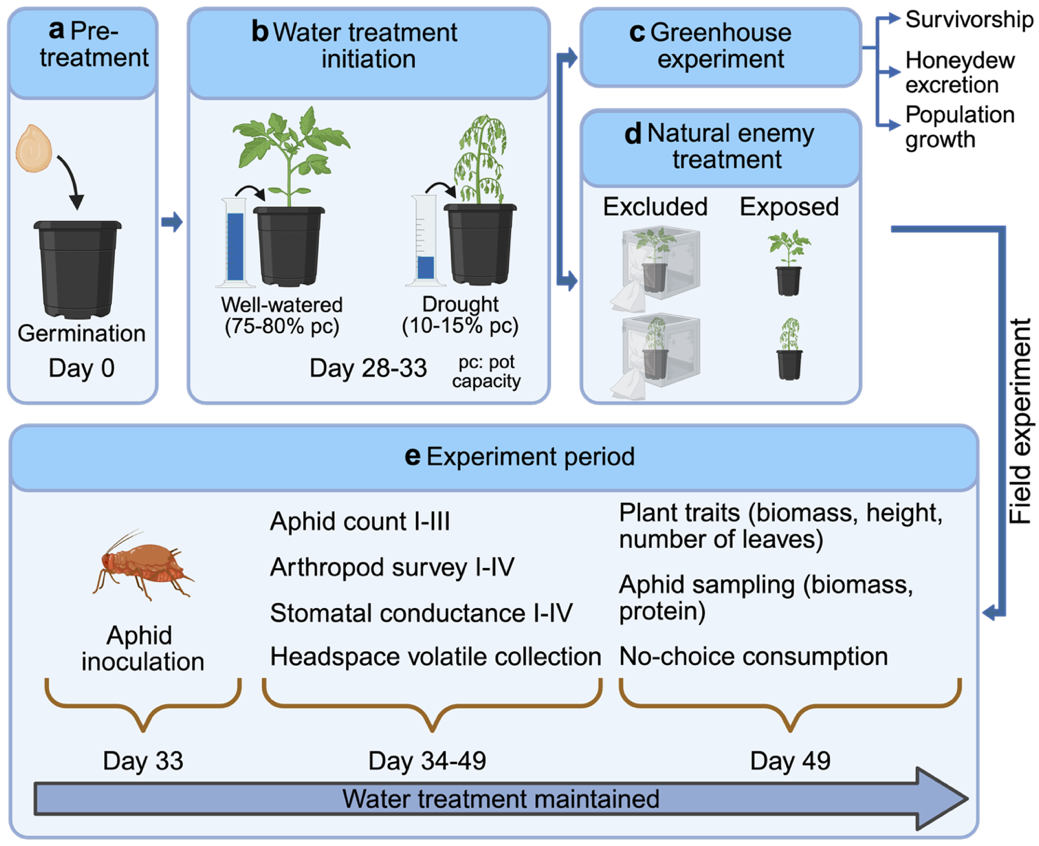

**Supplementary Fig. S1** Overview of the experimental design and timeline for both greenhouse and field experiments. (a) *Pre-treatment phase:* Seeds were germinated under optimal watering conditions beginning on Day 0. (b) *Water treatment initiation:* Between Days 28-33, plants were assigned to one of two soil moisture regimes, well-watered (75-80% pot capacity (pc) or drought (10-15% pc). (c) *Greenhouse experiment:* Plants from each water treatment were used to assess aphid performance under controlled conditions, including measurements of aphid survivorship, honeydew excretion, and population growth. (d) *Natural enemy treatment (Field experiment):* Concurrently, plants were transferred to field cages and subjected to natural enemy *exclusion* or *exposure* treatments to examine top-down effects on aphid populations. (e) *Field experiment period:* On Day 33, aphids were inoculated onto plants. From Days 34-49, aphid counts (I-III), arthropod surveys (I-IV), stomatal conductance measurements (I-IV), headspace induced plant volatile collections were conducted. Final measurements on Day 49 included plant traits (biomass, height, and leaf number), aphid biomass and protein content, and a no-choice ladybug consumption assay. Water treatments were maintained throughout both the greenhouse and field experiments.

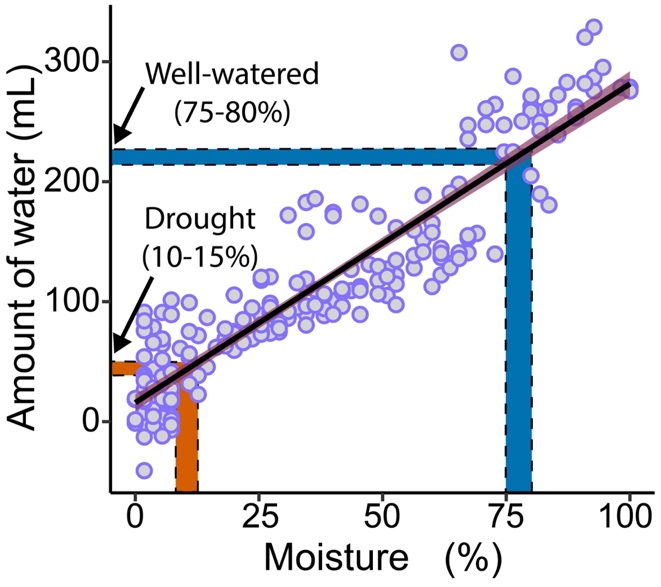

**Supplementary Fig. S2** Calibration curve illustrating the relationship between soil moisture percentage and water volume added to pots during watering treatments. This figure depicts the linear relationship between gravimetrically measured soil moisture (%) and the amount of water (mL) added to achieve those moisture levels in pots filled with potting mix. Moisture (%) was calculated daily from two drying trials using the equation: Moisture (%) = (Wet pot weight – Dry soil weight) / Water weight at full saturation × 100, where the average dry soil mass for a standard 4-inch² pot was 116.01 g (excluding pot weight). Full saturation was established by saturating soil and allowing it to drain for 24 hours to determine 100% pot capacity. Moisture was tracked until complete drying (0%).

The plotted regression line (black) follows the equation: Water added (mL) = 2.54 × Moisture (%) + 17.92. This equation was used to determine the volume of water required to reach target moisture levels. Blue and orange bars highlight the watering thresholds used in the experiment:

- Well-watered: 75–80% pot capacity (~208–221 mL)
- Drought: 10–15% pot capacity (~43–56 mL)

Water was applied approximately every day, but frequency was adjusted based on visual inspection of soil moisture. If pots appeared overly dry or saturated, the amount and timing of watering were modified based on estimated gravimetric moisture. For larger pots, the target water volumes were scaled proportionally to their dry soil content, based on the ratio of dry soil mass relative to small pots (e.g., ~660 g vs. ~180 g). The potting mix used in this experiment resembles coarse-textured soils and approximates a permanent wilting point at ~10% volumetric water content (v/v). This threshold informed the lower moisture limit for drought conditions.

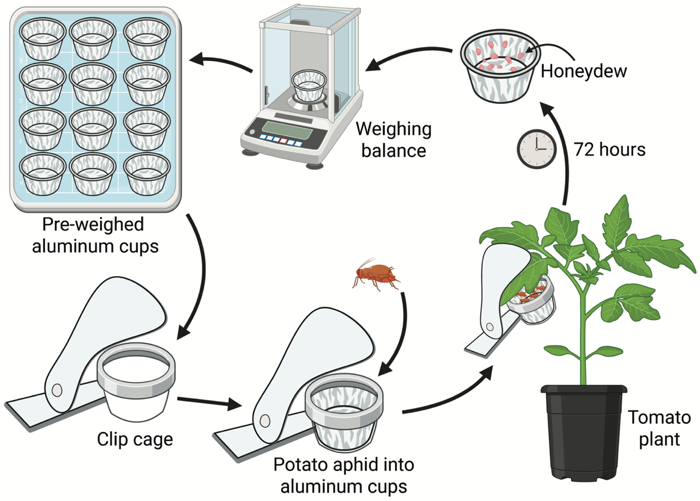

**Supplementary Fig. S3** Workflow illustrating the experimental procedure for quantifying aphid honeydew excretion. Pre-weighed aluminum foil cups were prepared and placed inside custom-designed clip cages. Ten apterous adult potato aphids were introduced into each cup, and the cages were secured onto tomato plants, targeting the abaxial leaf surface where aphids preferentially feed. After 72 hours, the clip cages were removed, and the aluminum cups were reweighed using a precision balance. Honeydew production was calculated as the change in cup mass before and after exposure and normalized per aphid (mg/aphid) based on final aphid counts. Only honeydew captured in the cups was quantified, as excretions typically fall away from the plant.

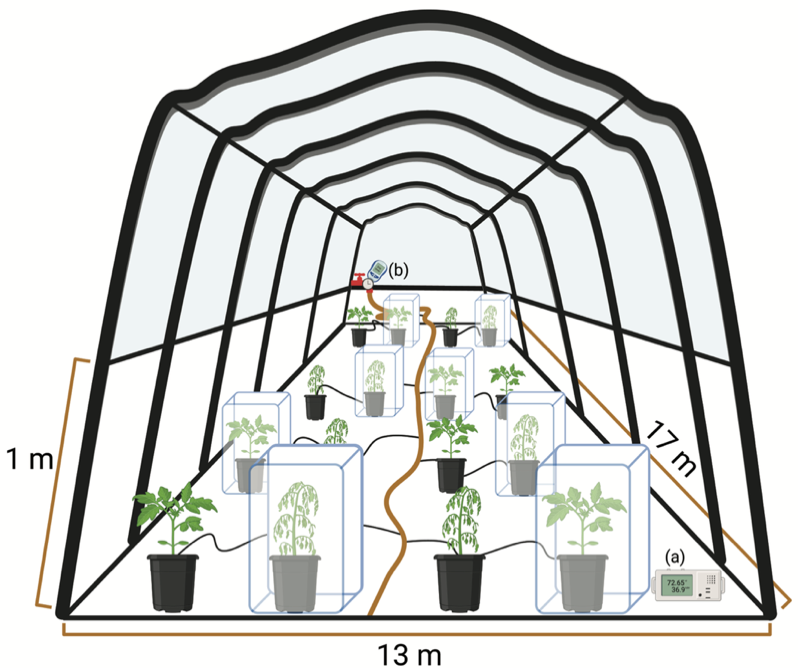

**Supplementary Fig. S4** Experimental layout within the rain-exclusion structure. The experiments were conducted from May to August at the Russell E. Larson Agricultural Research Center, Pennsylvania State University, USA. Plants were arranged inside a (13 × 17) m^2^ rain-exclusion structure with a transparent tarp roof and retractable sidewalls. The tarp was deployed to prevent rainfall, while a 1-meter open gap along each side allowed access by natural arthropod communities, including predators and parasitoids. Data loggers (a) were placed inside the structure to record environmental conditions. Plants were subjected to a two-way factorial design manipulating water availability and natural enemy access. Some plants were enclosed in mesh cages to exclude arthropods, while others remained uncaged to allow natural colonization. Drip irrigation lines provided water according to treatment, regulated by a watering timer (b).

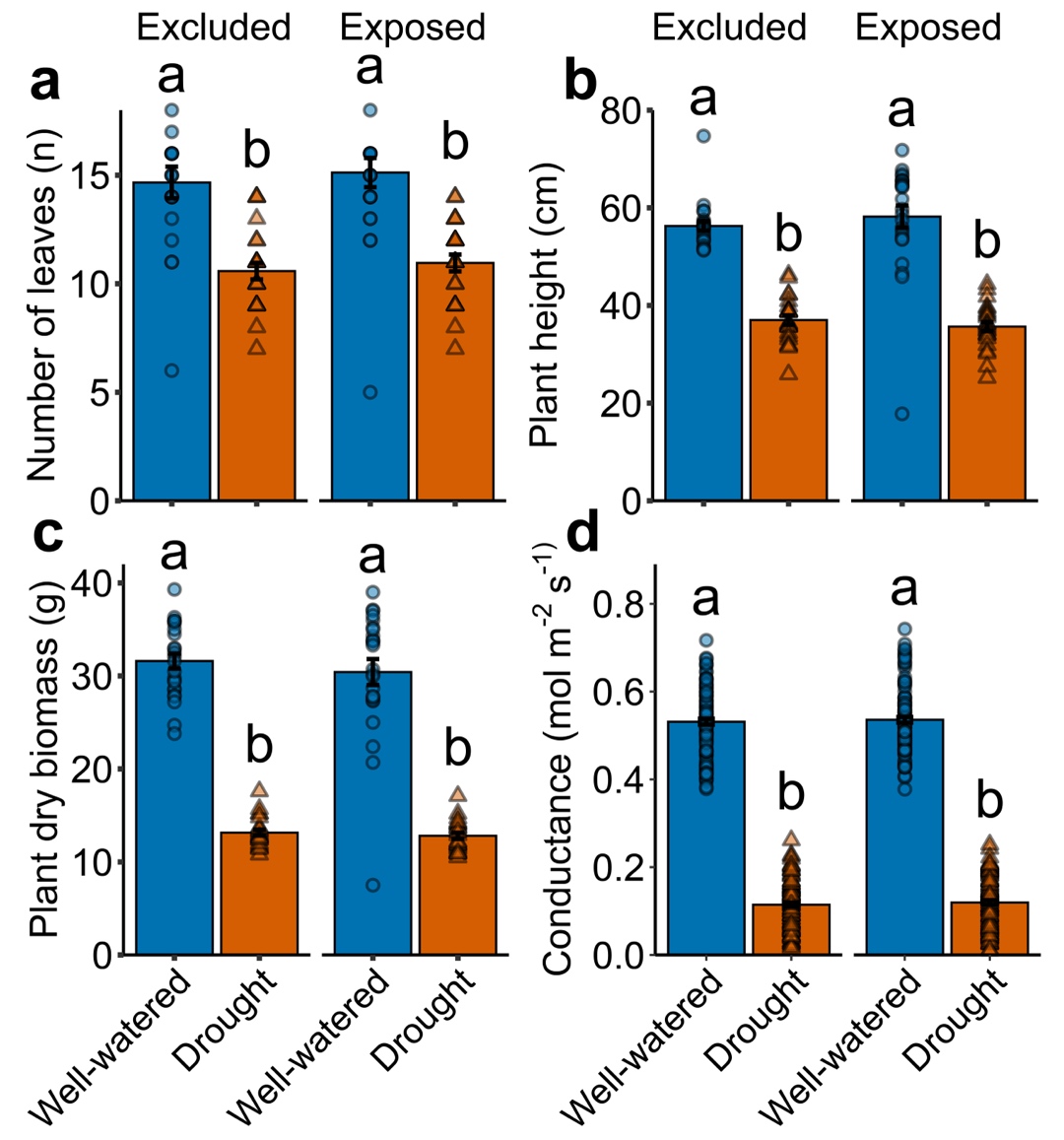

**Supplementary Fig. S5** Effect of water treatment (well-watered (blue bars) vs. drought (yellow bars)) and natural enemy exposure (excluded vs. exposed) on tomato plant traits. Bar plots show means (±1SE) for (a) number of leaves, (b) plant height, (c) plant dry biomass, and (d) stomatal conductance. Different letters above the bars indicate significant differences (*P*<0.05).

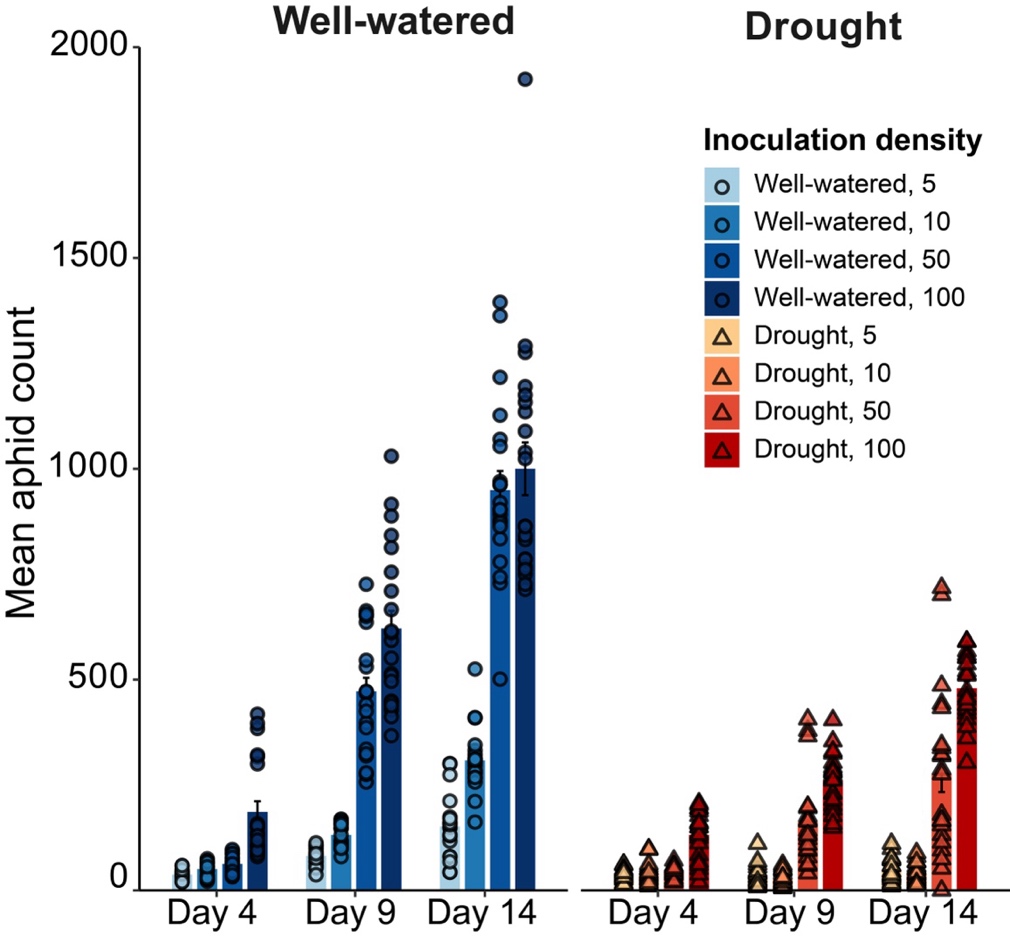

**Supplementary Fig. S6** Effects of drought stress on aphid population dynamics in greenhouse conditions. Mean aphid counts (± SE) over 14 days on tomato plants subjected to two water treatments (well-watered and drought-stressed) and four initial inoculation densities (5, 10, 50, and 100 aphids per plant). Bars represent mean aphid abundances at Days 4, 9, and 14 post-inoculation, with individual data points overlaid. Well-watered plants are shown in shades of blue and drought-stressed plants in shades of red, with color intensity corresponding to initial aphid density.

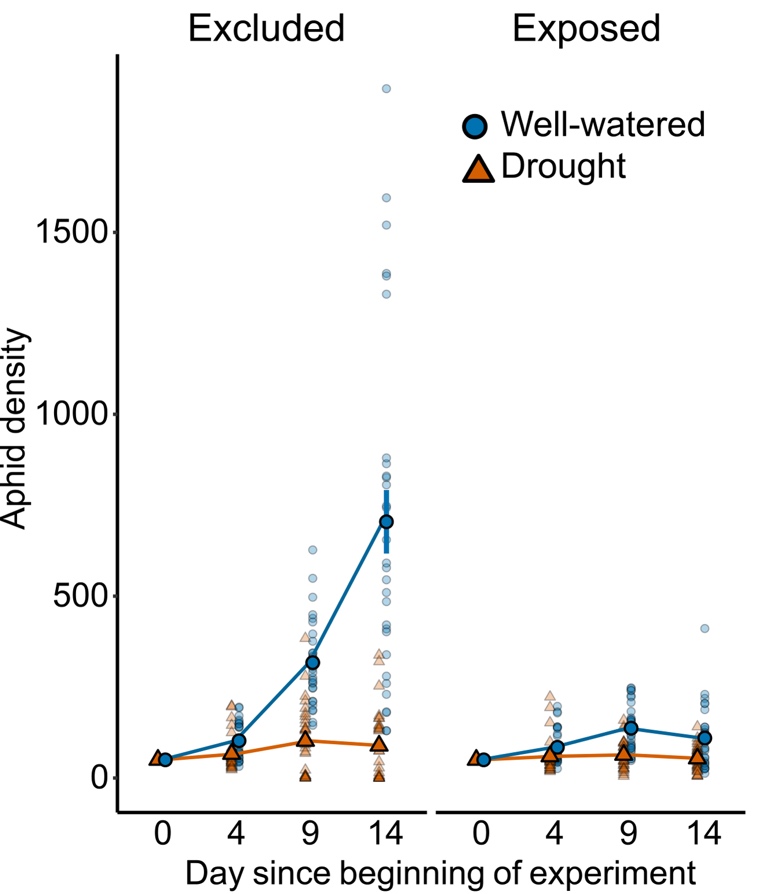

**Supplementary Fig. S7** Effects of drought stress and natural enemy exposure on aphid performance in field. Mean aphid counts (± SE) over 14 days on tomato plants subjected to water treatments (well-watered and drought-stressed) and natural enemy exposure (excluded and exposed). Point and error bar represent mean aphid abundances at Days 4, 9, and 14 post-inoculation, with individual data points overlaid. Well-watered plants are shown in shades of blue and drought-stressed plants in shades of orange.

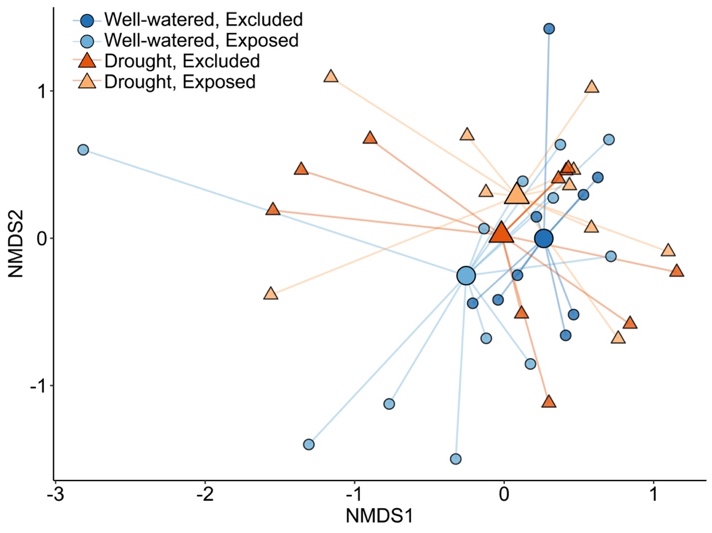

**Supplementary Fig. S8** Non-metric multidimensional scaling (*NMDS*) ordination depicting differences in herbivore-induced plant volatile (HIPV) profiles among tomato plants subjected to water availability (well-watered vs. drought-stressed) and natural enemy exposure (excluded vs. exposed) treatments. Each point represents the HIPV profile of an individual plant, with point shape indicating water treatment (circles: well-watered; triangles: drought-stressed) and fill indicating natural enemy treatment (dark blue or orange: excluded; light blue or orange: exposed). Centroids for each treatment group summarize the central tendency of volatile profiles. Differences in volatile composition across treatments were assessed via *PERMANOVA*, with significant effects of drought and natural enemy exposure on HIPV community structure.

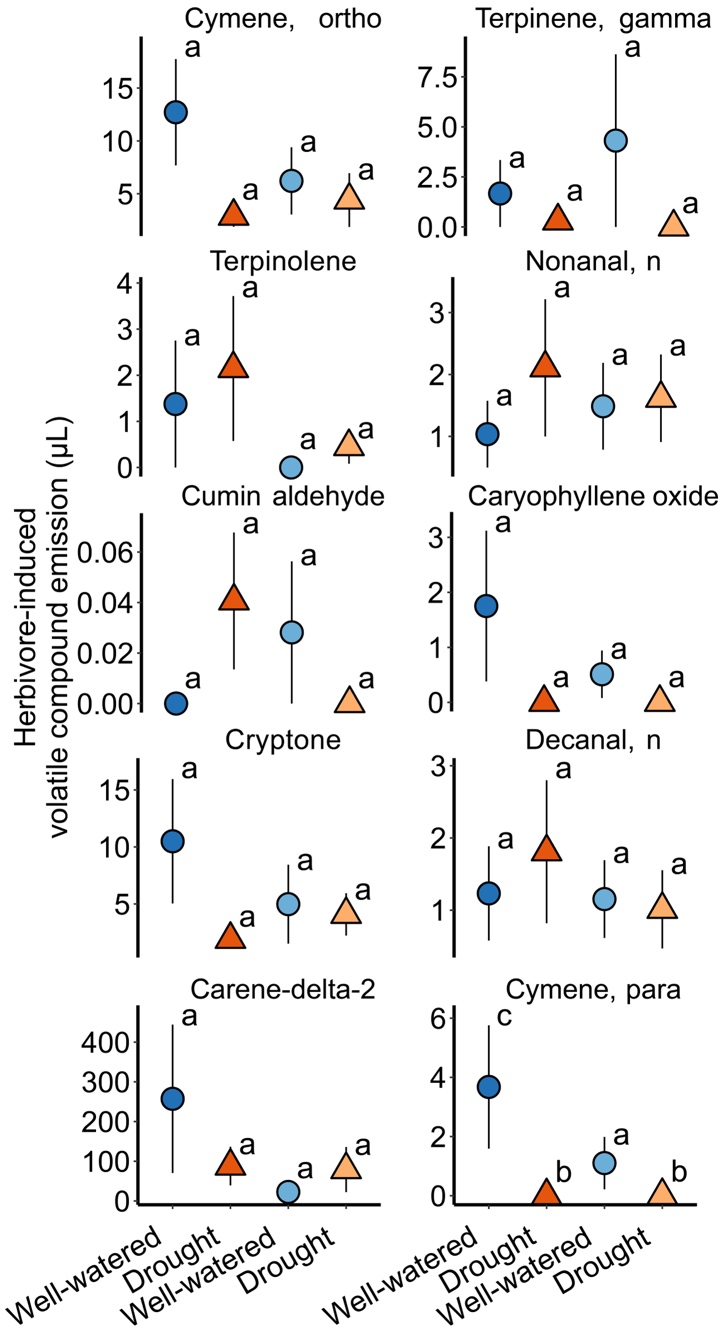

**Supplementary Fig. S9** Emission rates of individual herbivore-induced volatile organic compounds across water and natural enemy exposure treatments. Each panel displays the mean (± SE) emission of a single compound under four treatment combinations: well-watered vs. drought-stressed plants, either exposed to or excluded from natural enemies. Circles represent well-watered plants, triangles represent drought-stressed plants. Letters denote significant differences among treatment groups based on Tukey’s HSD tests (α=0.05). See Supplementary Table S4 for full statistical results.

**Supplementary Table S1** Results of likelihood ratio tests examining the effects of natural enemy exposure and drought-by-natural enemy interactions on plant traits.

| **Plant trait** | **Predictors** | **Chi-squared (χ²)** | **df** | **P-value** |
| --- | --- | --- | --- | --- |
| Leaf number | Natural enemy | 0.52 | 1 | 0.472 |
|  | Interaction (Drought × Natural enemy) | 0.002 | 1 | 0.960 |
| Plant height | Natural enemy | 0.04 | 1 | 0.840 |
|  | Interaction (Drought × Natural enemy) | 1.419 | 1 | 0.233 |
| Dry biomass | Natural enemy | 0.86 | 1 | 0.355 |
|  | Interaction (Drought × Natural enemy) | 0.255 | 1 | 0.613 |
| Stomatal conductance | Natural enemy | 0.22 | 1 | 0.642 |
|  | Interaction (Drought × Natural enemy) | 0.0013 | 1 | 0.971 |

**Supplementary Table S2** Effects of drought on arthropod community structure across feeding guilds and diversity metrics. Arthropod responses to water availability (drought vs. well-watered) using generalized linear mixed models (GLMMs), with Chi-square tests (*χ²*) used to evaluate fixed effects of water treatment on abundance, species richness, and Shannon diversity. Models were fit separately for each functional group and community-level metric. The table reports test statistics (Chi-square), degrees of freedom (*df*), and *P*-values. Statistically significant results (*P*<0.05) are bolded in the main text. Positive values indicate an increase under drought; negative values indicate a decline.

| **Metric** | **Guild** | **Chi-squared (χ²)** | **df** | **P-value** |
| --- | --- | --- | --- | --- |
| Abundance | Arthropod | 124.7360 | 1 | **0.000** |
|  | Chewer | 7.6530 | 1 | **0.006** |
|  | Omnivore | 2.8830 | 1 | 0.090 |
|  | Parasitoid | 2.7040 | 1 | 0.100 |
|  | Pollinator | 0.3070 | 1 | 0.580 |
|  | Predator | 90.6100 | 1 | **0.000** |
|  | Sap sucker | 36.0820 | 1 | **0.000** |
| Richness | Arthropod | 44.8540 | 1 | **0.000** |
|  | Chewer | 0.6190 | 1 | 0.432 |
|  | Omnivore | 0.7890 | 1 | 0.374 |
|  | Parasitoid | 3.3380 | 1 | 0.068 |
|  | Pollinator | 0.6370 | 1 | 0.425 |
|  | Predator | 27.6320 | 1 | **0.000** |
|  | Sap sucker | 24.5920 | 1 | **0.000** |
| Shannon diversity | Arthropod | 25.8620 | 1 | **0.000** |
|  | Chewer | 0.9580 | 1 | 0.328 |
|  | Omnivore | 2.2090 | 1 | 0.137 |
|  | Parasitoid | 2.3850 | 1 | 0.123 |
|  | Pollinator | 1.1960 | 1 | 0.274 |
|  | Predator | 40.0980 | 1 | **0.000** |
|  | Sap sucker | 16.6190 | 1 | **0.000** |

**Supplementary Table S3** Univariate ANOVA results for each compound testing the effects of Water treatment, Natural enemy caging status, and their interaction. Displayed are the degrees of freedom (df), F-values, and associated *P*-values for each effect.

| **Compound** | **Effect** | **df** | **F value** | **P value** |
| --- | --- | --- | --- | --- |
| Nonanal-n | Water | 1 | 0.508 | 0.480 |
|  | Natural enemy | 1 | 0.000 | 0.986 |
|  | Water: Natural enemy | 1 | 0.345 | 0.560 |
| Cryptone | Water | 1 | 1.757 | 0.193 |
|  | Natural enemy | 1 | 0.271 | 0.606 |
|  | Water: Natural enemy | 1 | 1.360 | 0.251 |
| Decanal-n | Water | 1 | 0.103 | 0.750 |
|  | Natural enemy | 1 | 0.383 | 0.540 |
|  | Water: Natural enemy | 1 | 0.264 | 0.610 |
| Carene-delta-2 | Water | 1 | 0.198 | 0.659 |
|  | Natural enemy | 1 | 1.883 | 0.178 |
|  | Water: Natural enemy | 1 | 1.578 | 0.217 |
| Cymene, ortho | Water | 1 | 2.841 | 0.100 |
|  | Natural enemy | 1 | 0.632 | 0.432 |
|  | Water: Natural enemy | 1 | 1.574 | 0.217 |
| Cymene-para | Water | 1 | 4.417 | 0.042 |
|  | Natural enemy | 1 | 1.527 | 0.224 |
|  | Water: Natural enemy | 1 | 1.500 | 0.228 |
| Terpinolene | Water | 1 | 0.515 | 0.477 |
|  | Natural enemy | 1 | 2.374 | 0.132 |
|  | Water: Natural enemy | 1 | 0.025 | 0.875 |
| Terpinene, gamma | Water | 1 | 1.307 | 0.260 |
|  | Natural enemy | 1 | 0.200 | 0.657 |
|  | Water: Natural enemy | 1 | 0.305 | 0.584 |
| Cumin aldehyde | Water | 1 | 0.040 | 0.843 |
|  | Natural enemy | 1 | 0.073 | 0.789 |
|  | Water: Natural enemy | 1 | 2.600 | 0.115 |
| Caryophyllene oxide | Water | 1 | 2.579 | 0.117 |
|  | Natural enemy | 1 | 0.929 | 0.341 |
|  | Water: Natural enemy | 1 | 0.903 | 0.348 |
